# Characterization and Calibration of the iQID Digital Autoradiography System for Direct Quantitative Imaging of Beta-Emitters in Tissue Samples

**DOI:** 10.64898/2026.01.27.701920

**Authors:** Ohyun Kwon, Sean P. Jollota, Adedamola O. Adeniyi, Justin J. Jeffery, Joseph B. Schulz, Lauren E. Wehner, Malick Bio Idrissou, Eduardo Aluicio-Sarduy, Brian W. Miller, Denis E. Bergeron, Reinier Hernandez, Larry A. DeWerd, Bryan P. Bednarz

**Author notes:** Corresponding author Address: 1111 Highland Avenue, 7109, Madison, WI 53705.

## Abstract

Autoradiography provides microscale mapping of radionuclide distributions, a promising approach to complement nuclear medicine imaging for small-scale radiopharmaceutical therapy (RPT) research. However, quantitative protocols for β-emitters remain under-established compared to those for α-emitters. In this work, the ionizing-radiation quantum imaging detector (iQID) digital autoradiography system was characterized and calibrated specifically for the theranostic β-emitter ^177^Lu. Spatial resolution, detection efficiency, background and minimum detectable activity, and depth dependence were characterized and compared to Geant4 Monte Carlo simulations. A methodology for converting count rates to activity was established, yielding a high linear response (range from 0 to 300 Bq). To validate the system for realistic measurement scenarios, cross-modality benchmarking was performed using a custom stacked multi-layer virtual water phantom to compare iQID performance with preclinical µSPECT/CT. The iQID system demonstrated an effective spatial resolution of ∼43 µm for ^177^Lu and achieved total activity estimates of (0.194 ± 0.022) MBq, agreeing within 2% with the dispensed reference (0.197 ± 0.015) MBq. Crucially, iQID exhibited superior quantitative accuracy for small-scale features (0.8 mm to 2.5 mm diameters), resolving activity concentrations in regions where µSPECT/CT performance was severely limited by partial volume effects. This study establishes a validated framework for quantitative ^177^Lu digital autoradiography, laying the groundwork for accurate activity estimation in *ex vivo* tissue samples.

## INTRODUCTION

Autoradiography has served as an important molecular imaging tool, allowing for the localization of radioactivity in biological specimens with the use of X-ray film or photographic emulsions (1,2). The capabilities of this imaging technology have advanced since it was introduced over a century ago, including the development of whole-body autoradiography by Ullberg in the 1950s and the subsequent innovation of phosphor imaging by Luckey in 1975, which facilitated quantitative whole-body autoradiography (3,4). Historically, this method used radiolabeled compounds, commonly low energy β-emitters such as ^14^C or ^3^H, for precisely tracing the distribution, metabolism, and retention of drugs within tissues (5–7). This approach has played an important role in drug discovery and development by enabling the characterization of pharmaceutical entities across diverse cellular and multicellular environments (1,8). A key advantage of autoradiography is its superior spatial resolution compared to typical *in vivo* nuclear medicine imaging modalities (9), such as preclinical small-animal μPET (∼1 mm) or μSPECT imaging (at best a few hundred micrometers).

However, traditional film-based or photographic emulsion autoradiography has several shortcomings that constrain its utility as a practical quantitative imaging technique. Early methods required long exposure times, typically ranging from days or weeks to months with long-lived radionuclides, which are inappropriate for *in vivo* studies (2). These techniques also suffered from a low dynamic range and susceptibility to image saturation, making it difficult to simultaneously visualize both high and low activity regions within a single sample (9). Furthermore, they were often only semi-quantitative, limited by a lack of linearity in film and by nonuniform thickness of photographic emulsions (1).

The introduction of digital autoradiography approaches mitigated many of these limitations. Applied in small-scale radiopharmaceutical therapy (RPT) research (10), digital autoradiography enables high-resolution activity mapping in thin tissue sections at defined time points and supports quantitative assessment of microscale radionuclide distributions across tumor and normal tissues. For example, Bäck and Jacobsson (2010) employed a digital α-particle camera to image the *ex vivo* distribution of ^211^At, achieving approximately 35 μm spatial resolution and generating quantitative activity patterns for different radiolabeled antibodies in tumor and normal tissues (11). Recent advancements in single-particle detection include the Timepix3-based Quantitative Particle Identification (QPID) system for spectral autoradiography (12) and the ionizing-radiation quantum imaging detector (iQID) (13).^1^ The iQID captures individual charged particle or photon emission events from a sample on an event-by-event basis, using a thin scintillator coupled to a high-sensitivity image sensor. Each decay event produces a light flash in the scintillator that is optically recorded and localized with sub-pixel precision, allowing accumulation in real time of a digital image in which each count corresponds to a detected decay and can be used to estimate activity concentration in the sample.

By providing accurate distributions of radionuclides at a near cellular level (∼tens of micrometers), digital autoradiography can support interpretation of therapeutic effect patterns and validation of biological models (14–16). A growing body of iQID studies with α-emitters has demonstrated these capabilities. Peter et al. reported small-scale dosimetry methods in two-dimensional (2D) canine biopsy sections and three-dimensional (3D) murine *ex vivo* tissues by pairing dose kernel-based absorbed dose rate maps with histological analysis (17,18). Similarly, Sahota et al. integrated iQID with coregistered morphology to analyze murine renal uptake patterns and facilitate close correspondence between absorbed dose rate maps and histology images (19). These examples indicate that advanced digital autoradiographic imaging provides the spatial resolution and quantitative performance needed to resolve nonuniform activity distributions in tissue samples, and they serve as a useful complement to nuclear medicine imaging modalities such as μSPECT/CT for activity quantification and method validation.

In this study, we focus on the characterization and calibration of the iQID digital autoradiography system for direct quantitative imaging of the theranostic β-emitter, ^177^Lu. Building on previous iQID studies with α-emitters and small-scale dosimetry, we established a practical methodology for measuring β-emitter activity distributions using surface-dispensed sources on microscope glass plates as a preparatory step toward *ex vivo* tissue measurements, with validated accuracy and precision. We evaluated detector performance, characterized various metrics, and calibrated the iQID camera with ^177^Lu. Finally, we performed a cross-modality benchmark using a custom stacked multi-layer (SML) phantom constructed from virtual water.

Prepared by patterned dispensing and drying of a radionuclide solution, the phantom enabled a direct comparison of images acquired with µSPECT/CT and iQID. This benchmarking approach offers a validation method that is challenging to achieve with α-emitters, which often lack imageable γ emissions for µSPECT imaging. The following sections describe our iQID system calibration procedures and results demonstrating its utility for high-resolution quantitative autoradiography.

## MATERIALS AND METHODS

### iQID camera and scintillating phosphor screens

The iQID camera (QScint Imaging Solutions, LLC, Tucson, AZ) is a digital autoradiography system that utilizes electro-optical gain to amplify scintillation light from a phosphor screen, enabling single event detection with a CCD/CMOS sensor (20). The system includes a 40 mm diameter image intensifier that is optically coupled to a 2448 × 2048 pixel camera that operates at a maximum frame rate of 75 s^−1^ (full frame). The image intensifier amplifies scintillation light from the phosphor screen upon radiation interaction, and the coupling lens projects the intensified image onto the sensor, where radiation events appear as localized flashes of light forming event clusters (13).

Different scintillating phosphor screens can be selected based on particle type and energy sensitivity. A silver-doped zinc sulfide (ZnS:Ag) screen with a 25 µm phosphor layer on a 50 µm to 100 µm transparent polyester substrate was used for α-emitting radionuclides, while a gadolinium oxysulfide (Gd₂O₂S, Gadox) screen with a 40 µm phosphor layer on a 100 µm polyester substrate was used for α- and β/γ-emitters. The properties of the two scintillating phosphor screens are summarized in **Table 1**.

**Table 1.**
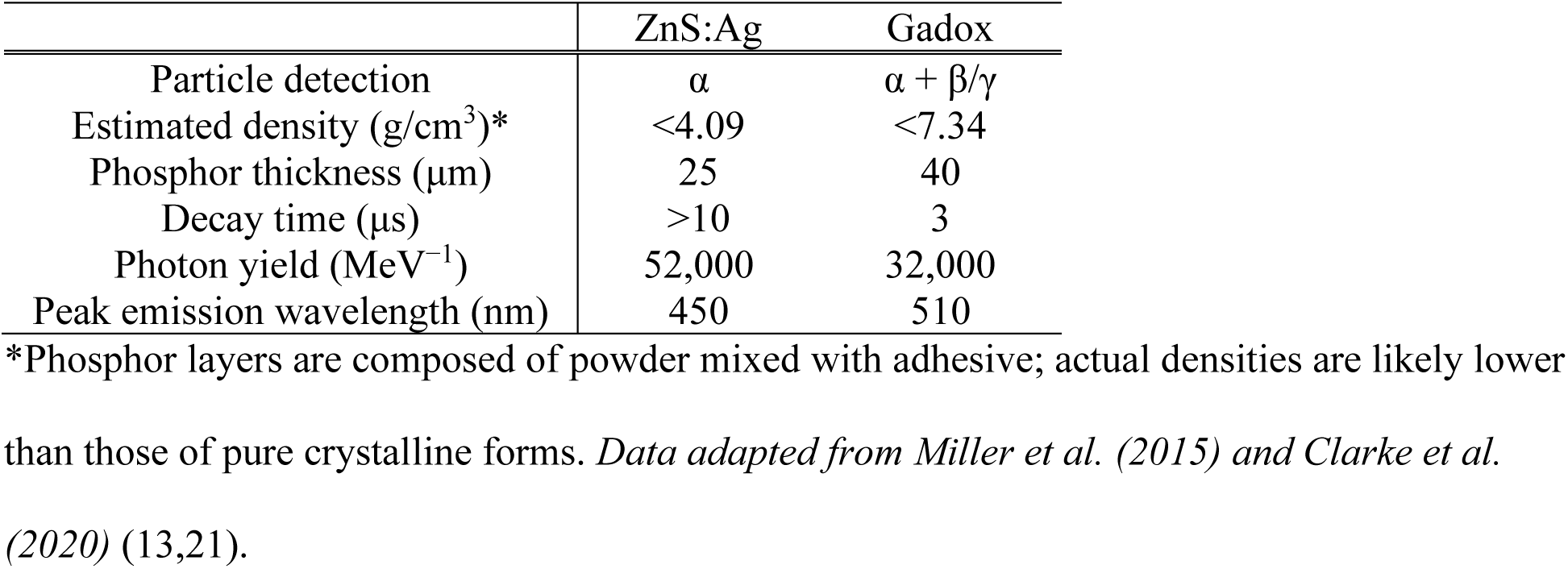
Properties of phosphor screens.

|  | ZnS:Ag | Gadox |
| --- | --- | --- |
| Particle detection | $\alpha$ | $\alpha + \beta/\gamma$ |
| Estimated density ($\text{g}/\text{cm}^3$)* | <4.09 | <7.34 |
| Phosphor thickness ($\mu\text{m}$) | 25 | 40 |
| Decay time ($\mu\text{s}$) | >10 | 3 |
| Photon yield ($\text{MeV}^{-1}$) | 52,000 | 32,000 |
| Peak emission wavelength (nm) | 450 | 510 |

\*Phosphor layers are composed of powder mixed with adhesive; actual densities are likely lower than those of pure crystalline forms. *Data adapted from Miller et al. (2015) and Clarke et al. (2020) (13,21).*

### Preparation of radioactive sample and activity validation

^177^LuCl_3_ solution was obtained from SHINE Technologies (Janesville, WI) in HCl solution and diluted as required for experimental use. Working solutions were prepared in phosphate-buffered saline (PBS), then dispensed by micropipette onto the corresponding plates or phantoms and dried at room temperature overnight prior to imaging. Activity concentrations were determined using a Capintec CRC-55tW dose calibrator (Mirion Technologies, Atlanta, GA). Solutions were transferred into 1.5 mL Eppendorf tubes using a micropipette and the calibrator was applied with manufacturer calibration. For the cross-modality benchmark, selected preparations were measured in the Eppendorf tubes using a high-purity germanium (HPGe) detector at the University of Wisconsin Cyclotron Laboratory. These reference measurements relied on an efficiency calibration curve established using a mixed γ standard at a fixed source-to-detector distance. All activity measurements were performed using fixed, reproducible geometry consistent with the calibrated instrument records. Temporal decay corrections for all datasets were standardized using a half-life of (6.647 ± 0.004) days (22), introducing negligible uncertainty (< 0.1%) to the final activity estimates.

### Characterization of iQID

#### Spatial resolution

The iQID camera has an intrinsic spatial resolution of 19.53 μm (13). As β particles emitted from ^177^Lu (*E*_max_ = 496.8 keV (23)) have ranges in water of approximately 0.62 mm (24), particle transport is expected to broaden the point response beyond the pixel pitch. To quantify this effect, we performed Geant4 Monte Carlo simulations (version 11.1.0) (25–27) using the G4RadioactiveDecay library derived from the Evaluated Nuclear Structure Data Files (ENSDF) and the experimental geometry (microscope glass plate, 2D point source, mylar sheet, Gadox phosphor screen, and iQID faceplate). Electron transport was tracked through all layers, and energy deposition in the Gadox was tallied in 19.53 × 19.53 µm^2^ bins to construct a simulated point-spread function (PSF). The full width at half maximum (FWHM) of the PSF was then computed. All simulations were run for at least 10^7^ primary ^177^Lu decays unless otherwise specified, ensuring statistical convergence and negligible uncertainty. Based on previous studies (9), a 5 keV energy threshold for iQID event detection was adopted and applied in the simulations. Experimentally, the detection threshold was tuned to balance retention of higher energy β events against suppression of low-level background.

#### Detection efficiency

The detection efficiency of the iQID varies with particle type, energy, and imaging settings. Previous studies have reported near 100% detection efficiency for α particles in a 2π geometry (13,21). In contrast, the detection efficiency for β particles has not been established experimentally and is expected to vary due to different factors such as scattering, secondary electron contributions, and pile-up effects. To estimate the absolute detection efficiency for ^177^Lu, Geant4 simulations were performed using the experimental geometry and the 5 keV energy threshold.

#### Background and minimum detectable activity

Intrinsic system and environmental background signals were measured under the same experimental imaging conditions in the absence of radioactive sources. To distinguish true particle signals from background noise (e.g., electronic noise or ambient radiation), the intensifier input voltages were optimized for α- and β-emitting radionuclides to minimize false pixel signals while remaining below the raw saturation threshold. The theoretical minimum detectable activity (MDA) for ^177^Lu was calculated based on the Currie equation (28) using simulation-based absolute detection efficiencies, as shown in Equation (1).

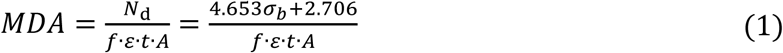

where *N*_*d*_ is the critical level of detection (in counts) required to ensure a false-negative rate no greater than 5%, and σ_*b*_ is the standard deviation of the background counts (*N*_*b*_), defined as σ_*b*_ = √*N*_*b*_ for Poisson statistics. Additionally, *f* is the branching ratio of the radionuclide (emitted particles per decay), ε is the absolute detection efficiency (detected counts per emitted particle), *t* is the acquisition time (s), and *A* is the effective detection area (cm^2^).

#### Depth dependency

Depth-dependent response was evaluated by placing stacks of mylar sheets between a dispensed 2D circular ^177^Lu source and the Gadox phosphor screen. The required thicknesses of mylar stacks were selected to reduce count rates from 100% to approximately 10% level, based on emitted particle energies from the International Atomic Energy Agency (IAEA) Nuclear Data Services (NDS) Live Chart of Nuclides (29) and the continuous slowing down approximation (CSDA) ranges and stopping-power values from the National Institute of Standards and Technology (NIST) ESTAR database (30). Geant4 simulations were performed with the experimental geometry, the 5 keV energy threshold, and the same thickness configurations for direct comparison with measurements.

#### Calibration of iQID

Calibration measurements were performed under conditions matching the preclinical imaging setup. To create a calibration source, ^177^Lu solution was dispensed onto 2.5 cm × 7.5 cm × 0.1 cm Superfrost Plus microscope glass plates (Fisher Scientific, Pittsburgh, PA). A single mylar sheet was then adhered to the plate by taping to maintain consistency, prevent contamination, and avoid activity loss across multiple measurements. This source was placed in a custom-designed holder to ensure reproducible positioning relative to the phosphor screen and the iQID camera.

Using the optimal detector settings established during characterization, a count rate to activity (s^−1^ to Bq) calibration curve was generated for ^177^Lu. For this procedure, the same source was measured repeatedly over time to leverage physical radioactive decay, thereby avoiding inter-source variability. The net count rate was calculated by subtracting the measured background count rate. A linear calibration model relating net count rate to activity was fit with the intercept constrained to zero, consistent with the expectation that zero activity yields zero net count rate under fixed imaging conditions. The calibration activity range was selected based on expected levels observed in previous preclinical studies.

### Cross-modality multi-scale phantom benchmarking

#### Phantom design and source preparation

To enable cross-modality validation of calibrated iQID imaging against μSPECT/CT (U-SPECT, MILabs, Utrecht, the Netherlands), a Derenzo phantom (D270825-GrIT, Phantech Co.) was utilized as a comparative benchmark for μSPECT/CT imaging. This reference phantom contained 10 mm length rods with diameters of 0.8 mm, 1.0 mm, 1.25 mm, 1.5 mm, 2.0 mm, and 2.5 mm. Based on this reference pattern, a custom SML phantom was fabricated using virtual water (University of Wisconsin Medical Radiation Research Center (UWMRRC) and University of Wisconsin Machine Shop). This phantom consisted of ten virtual water layers (28 mm diameter, 1 mm thick), each machined with 0.1 mm depth well patterns matching the Derenzo rod diameters. **Figure 1** shows the pattern configuration and appearance of both phantoms.

**Figure 1.**
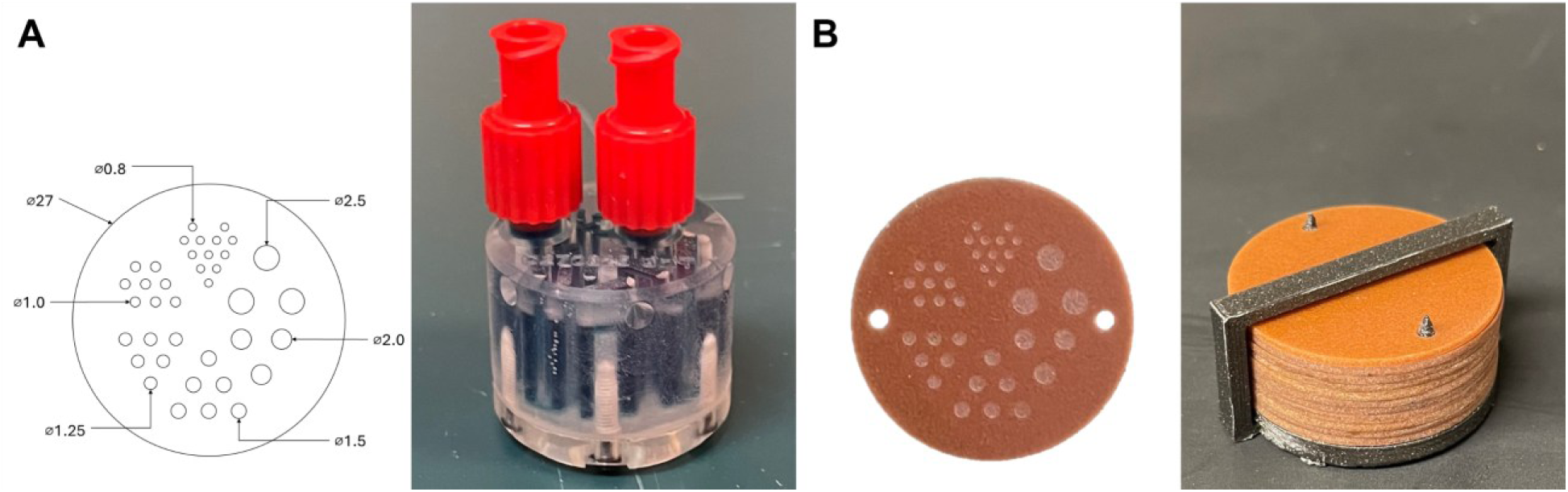
Phantom configurations. (A) Derenzo phantom containing six rod diameters (10 mm length). (B) SML phantom with virtual water layers (1 mm thick) containing identical well patterns (0.1 mm depth).

A nominal total activity of 0.2 MBq ^177^Lu was prepared for each phantom. For the Derenzo phantom, the ^177^Lu solution was injected and sealed. For the SML phantom, a fixed activity concentration ^177^Lu solution was dispensed into the wells and dried. A single mylar sheet was applied to each layer of the SML phantom to prevent contamination. The layers were then stacked using a custom 3D-printed polylactic acid (PLA) alignment frame (density ∼1.24 g·cm^−3^, *Z*_*eff*_ ≈ 7) to form a 10 mm length cylindrical phantom and covered with a 1 mm thick non-patterned layer.

Both the Derenzo and SML phantoms were individually imaged using μSPECT/CT (8-hour acquisitions per phantom). Five weeks later, the SML phantom was disassembled and each layer was imaged separately using the calibrated iQID system.

#### Recovery coefficients analysis

Quantitative imaging accuracy and resolution performance were assessed by comparing activity concentrations from μSPECT/CT and iQID images after decay correction to a common reference time (the μSPECT/CT scan time). Geometric correction factors derived from Geant4 simulations were applied to the iQID data to account for the 0.1 mm well depth relative to the flat calibration geometry. Regions of interest (ROIs) were defined based on μCT images for μSPECT and on shadowgraph images for iQID, where shadowgraph refers to CCD/CMOS images of the perforated well pattern layer acquired by recording light transmitted from a uniformly illuminated lid sheet in the absence of phosphor screens. Recovery coefficients (RCs) were calculated for the six feature diameters (0.8 mm to 2.5 mm) to quantify partial volume effects. The analysis compared the Derenzo phantom (μSPECT/CT image) and the SML phantom (μSPECT/CT and iQID images).

## RESULTS

### Characterization of iQID

#### Spatial resolution

The effect of β-particle transport on spatial resolution was quantified using Geant4 simulations. The resulting simulated PSF yielded a FWHM of 43.33 μm, which corresponds to 2.22 pixels. This simulated resolution is broader than the iQID camera’s intrinsic spatial resolution (19.53 μm), as anticipated due to the range of ^177^Lu β-particles. Figure 2 shows the simulation geometry in air (microscope glass plate, ^177^Lu source, a single 3 × 3 cm^2^ mylar sheet, a 5 × 5 cm^2^ phosphor screen, and the iQID image intensifier faceplate) and the results for the PSF with its FWHM analysis.

**Figure 2.**
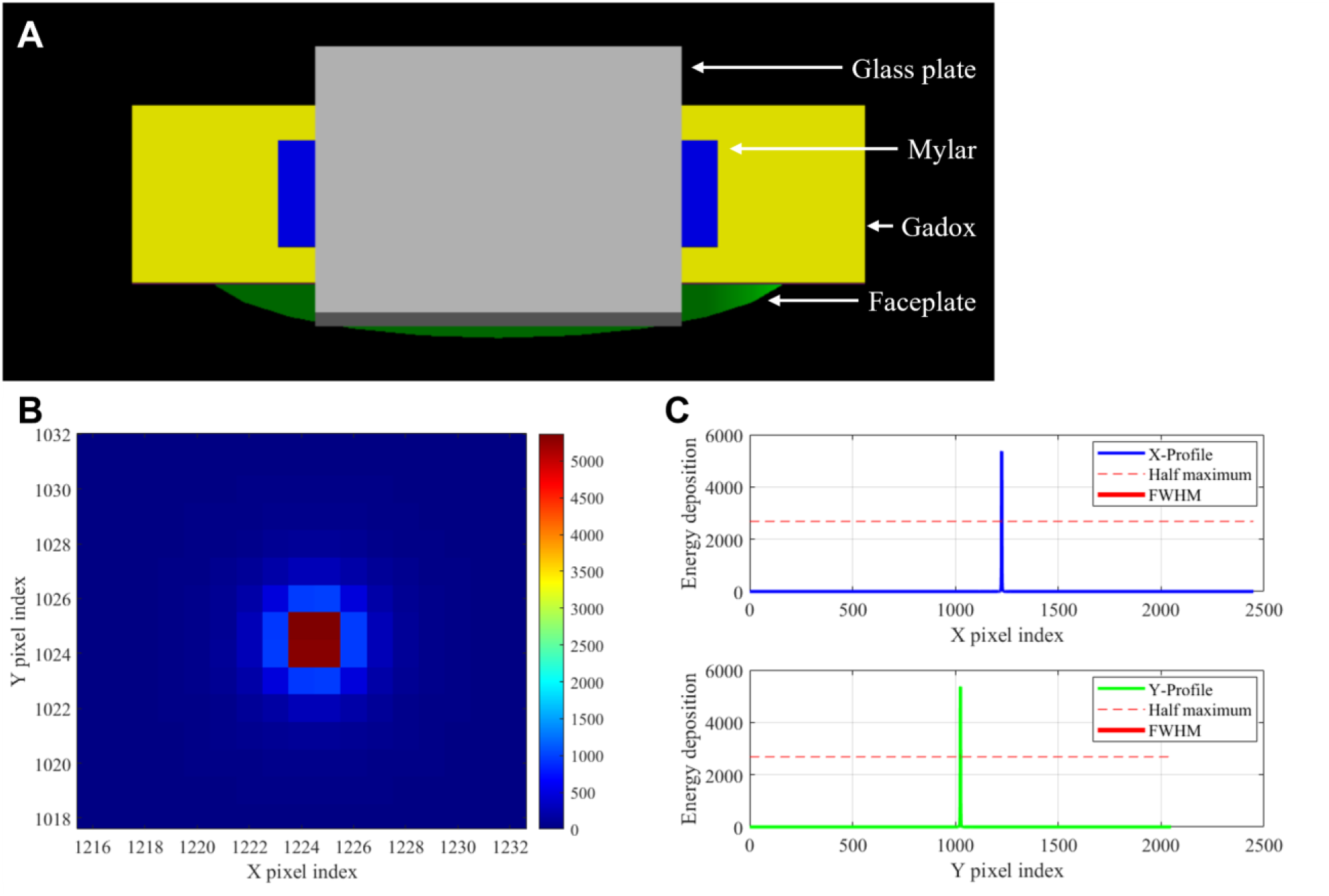
Geant4 simulation geometry and spatial resolution analysis. (A) 3D model of the experimental setup: microscope glass plate (gray), point source, mylar sheet (blue), Gadox screen (yellow), and iQID faceplate (green). (B) Heatmap of the simulated point spread function derived from β-particle energy deposition (MeV). (C) Horizontal and vertical profiles through the point spread function peak (in 2448 × 2048 pixel resolution) with the corresponding full width at half maximum.

#### Detection efficiency

Geant4 simulations were first validated for ^177^Lu by modeling a point source in a vacuum and comparing the emission results with the reference data of the IAEA NDS Live Chart of Nuclides. The average relative differences for β and γ emissions were 3.8%. Following validation, the 5 keV energy threshold was applied, and the full imaging geometry with a 5 mm diameter disc source was implemented in the simulation (Figure 2A). The number of scored events per decay reaching the phosphor layer was recorded. These simulation results were used to estimate the absolute detection efficiency of the iQID system for ^177^Lu with the Gadox scintillator configuration, as summarized in **Table 2**.

**Table 2.** Comparison of ^177^Lu emission yields (IAEA reference vs. Geant4 vacuum simulation) and simulated detection efficiency at the Gadox screen.

| Source | Method | Geometry | $\beta^-$ | $\gamma$ | Detection Efficiency |
| --- | --- | --- | --- | --- | --- |
| $^{177}\text{Lu}$ | IAEA | Vacuum | 0.62 | 0.13 | - |
|  | Geant4 | Vacuum | 0.60 | 0.13 | - |
|  |  | iQID system | 0.80 | 0.05* | 0.85 |

\*Photon was collected from collisions under the iQID system geometry.

#### Background and minimum detectable activity

Intrinsic background signals were measured over input voltages from 1.0 V to 5.0 V using maximum gain (47.99 dB), maximum shutter time (13.18 ms), and default detection thresholds (threshold = 100; cluster threshold = 1 pixel), as shown in Figure 3. Acquisition durations ranged from 2 hours to 15 hours for voltages ≤ 3.0 V, 5 minutes to 2 hours for 3.0 V to 4.0 V, and were reduced to < 5 minutes above 4.0 V to limit device load. Count rates ranged from (0.0030 ± 0.0004) s^−1^ to (36.3 ± 0.7) s^−1^ for electrical noise only (no phosphor screen), (0.0036 ± 0.0006) s^−1^ to (46.6 ± 0.9) s^−1^ for the ZnS:Ag screen, and (0.0029 ± 0.0001) s^−1^ to (50.4 ± 1.2) s^−1^ for the Gadox screen. To approximate typical signal levels expected from a ^177^Lu source, an input voltage of 3.0 V was selected as optimal for the Gadox screen. Scan time selection was guided by the coefficient of variance from the background noise at 3.0 V with the Gadox screen, which measured 3.06%, 1.58%, 1.03%, and 0.62% for durations of 10 minutes, 30 minutes, 1 hour, and 2 hours, respectively. Final background measurements at 3.0 V with the Gadox screen for β-emitting radionuclides (2 hours) yielded a count rate of (1.999 ± 0.065) s^−1^.

**Figure 3.**
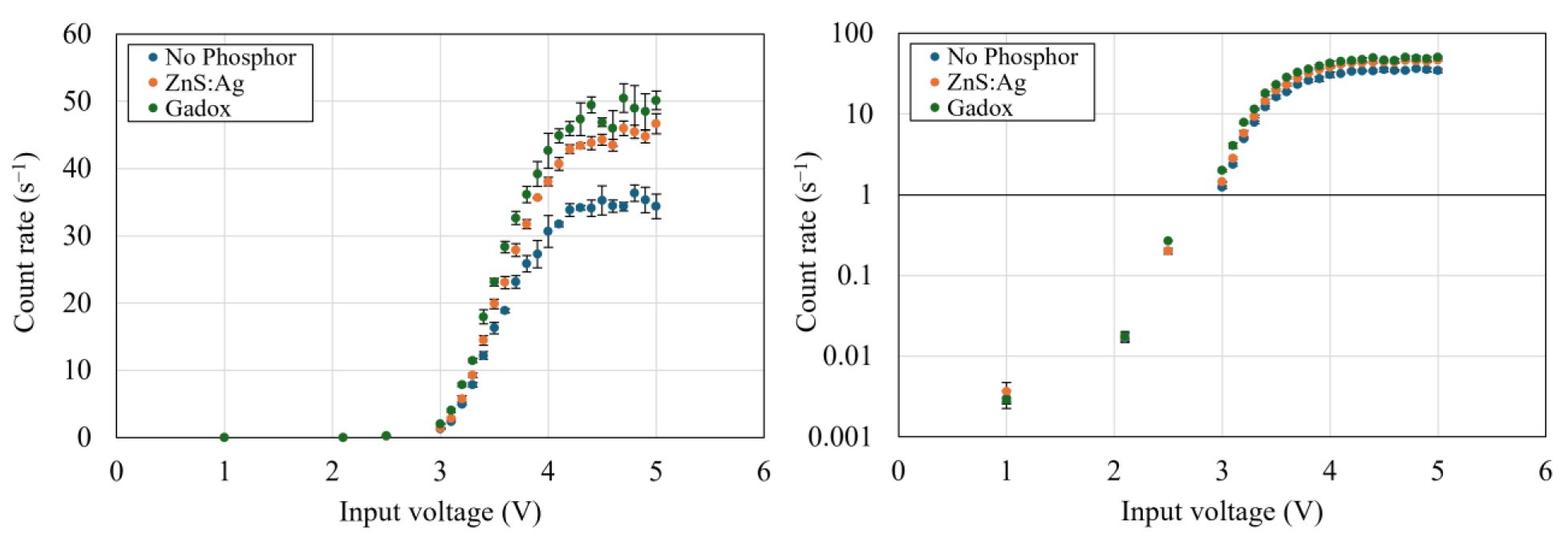
Background count rates as a function of input voltage plotted on linear (left) and logarithmic (right) scales. Data are shown for (1) electrical noise (no screen), (2) ZnS:Ag, and (3) Gadox screens, acquired at maximum gain and shutter settings.

Using this background value, the MDA for ^177^Lu at these settings was calculated based on the absolute detection efficiencies, summarized in **Table 3**.

**Table 3.** Calculated minimum detectable activity of the iQID camera for ^177^Lu.

| Source | Phosphor Screen | Background Count | Time (s) | $f$ | $\varepsilon$ | MDA (Bq/cm<sup>2</sup>) |
| --- | --- | --- | --- | --- | --- | --- |
| $^{177}\text{Lu}$ | Gadox | 14393 | 7200 | 1 | 0.85 | 1.15 |

#### Depth dependency

Experimental count rates from ^177^Lu measured through stacked mylar sheets were compared with Geant4 simulation results (relative counts per decay). Measurements used 1, 11, 26, 47, and 74 mylar sheets (3 μm, 33 μm, 78 μm, 141 μm, and 222 μm). The absolute count rate, decay-corrected to the first measurement time point, was obtained by subtracting background count rate from the acquired count rate and then extrapolated to 0 sheets using a bi-exponential fit (R^2^ = 0.9994) to estimate the relative count rate. The average absolute percent difference between the two datasets was 5.3%, calculated as the mean of the absolute percent difference (mean|(simulated − measured)/measured| × 100). A quantitative comparison of all measured and simulated results is provided in **Table 4**. Figure 4 shows simulated counts per decay versus β-particle energy, grouped by mylar thickness.

**Figure 4.**
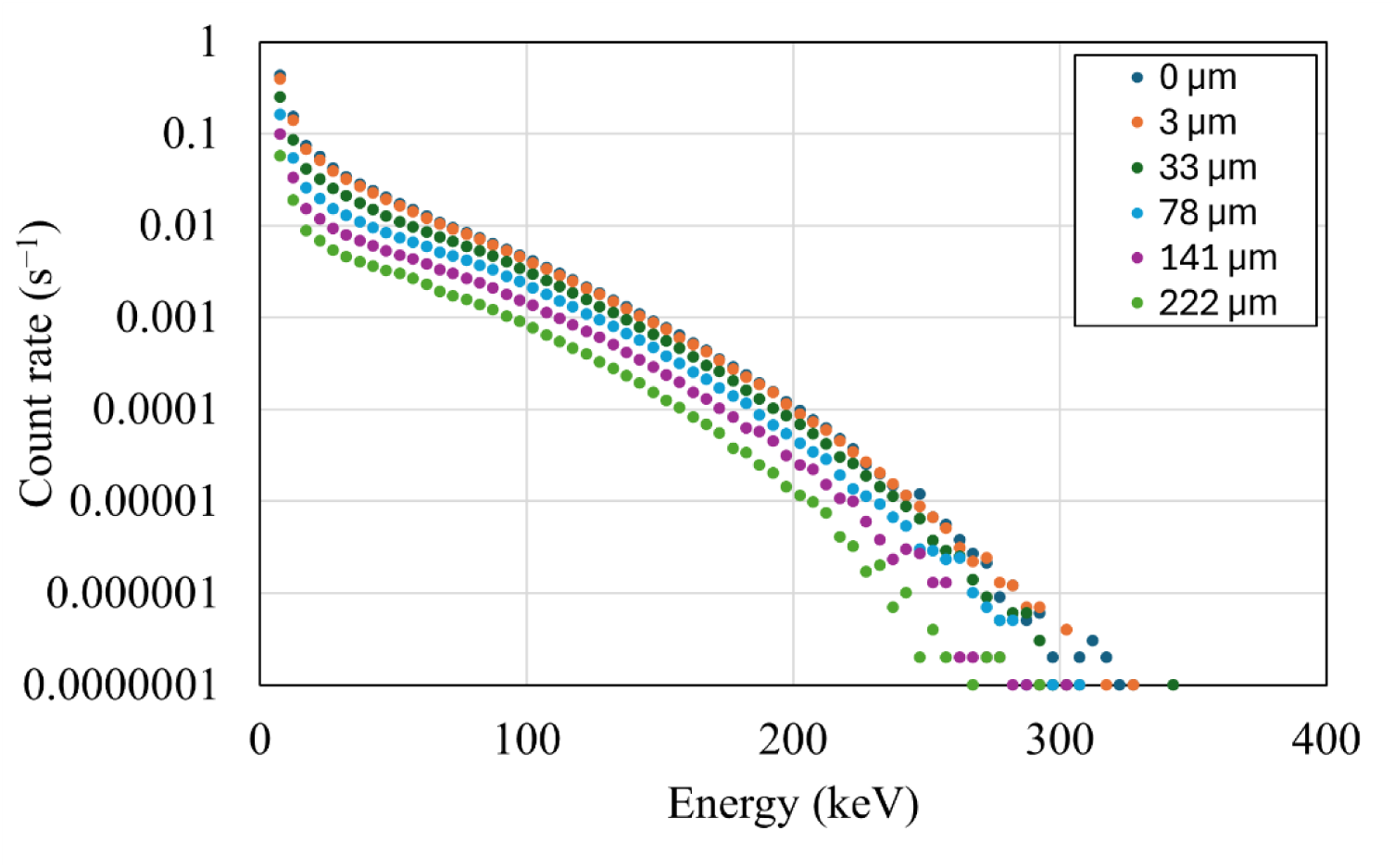
Simulated histograms for ^177^Lu β particles as a function of deposited energy, shown for varying mylar thicknesses and distance between source and phosphor screen.

**Table 4.** Comparison of measured (decay-corrected count rates) and simulated (counts per decay) ^177^Lu results across varying mylar thicknesses.

| Source | Phosphor Screen | Mylar | Thick ($\mu\text{m}$) | iQID Count Rates | | Geant4 Efficiencies | | %Diff. |
| --- | --- | --- | --- | --- | --- | --- | --- | --- |
| | | | | Abs. ($\text{s}^{-1}$) | Rel. | Abs. Entry (counts/decay) | Rel. Entry | |
| $^{177}\text{Lu}$ | Gadox | 0 | 0 | - | 1.000 | 0.860 | 1.000 | - |
|  |  | 1 | 3 | 196.7 | 0.949 | 0.799 | 0.928 | -2.2 |
|  |  | 11 | 33 | 124.6 | 0.601 | 0.532 | 0.618 | 2.9 |
|  |  | 26 | 78 | 80.9 | 0.390 | 0.351 | 0.407 | 4.4 |
|  |  | 47 | 141 | 48.4 | 0.234 | 0.214 | 0.248 | 6.4 |
|  |  | 74 | 222 | 32.1 | 0.155 | 0.119 | 0.138 | -10.9 |

#### Calibration of iQID

The ^177^Lu calibration curve was established using the optimized detector configurations identified during characterization (Gadox screen, 3.0 V input voltage, default internal settings). Utilizing the single-source decay method, seven calibration points covering 0 Bq to 300 Bq were acquired (2 hours per scan). Linear regression analysis yielded a calibration factor of 1.5126 Bq/(s^−1^) with a negligible intercept, confirming effective background subtraction. The regression showed high linearity (R^2^ = 0.9965), with minimal random uncertainty (Type A) in the fit. The cumulative systematic uncertainty (Type B) was estimated at 3.2%, dominated by the reference measurement accuracy of the Capintec CRC-55tW in standard tube geometry (±2.5%) (31) and volumetric pipetting precision (±2.0%) (32). The calibration results are summarized in Figure 5 and **Table 5**.

**Figure 5.**
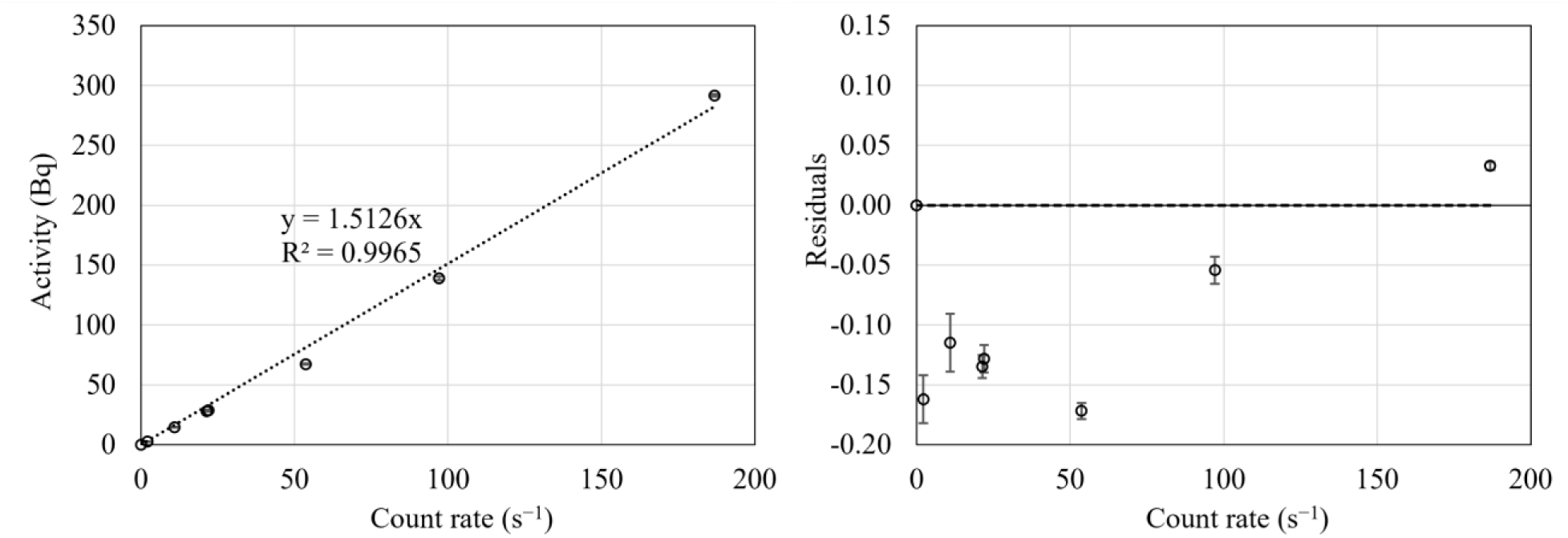
Calibration curve of ^177^Lu activity plotted against iQID count rate (left). The linear regression equation and R^2^ value are shown. Error bars indicate standard deviations of the background-corrected measurements. The corresponding relative residuals plotted against the same count rate (right), calculated as (measured – fitted) / fitted. Error bars represent the measurement uncertainty propagated relative to the fitted value.

**Table 5.**
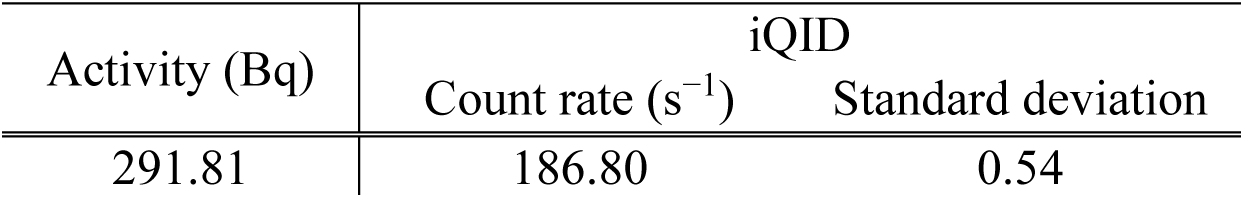
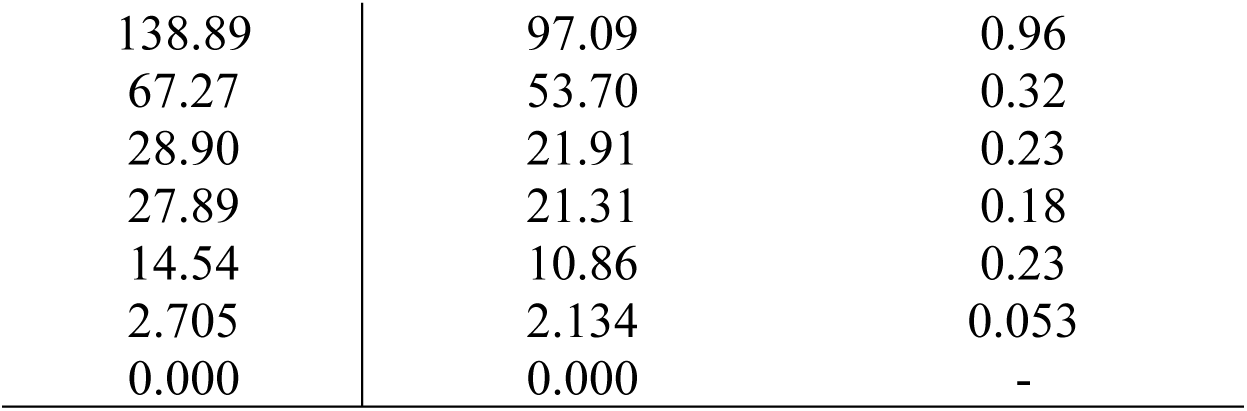
^177^Lu activity calibration using iQID measurements. Count rates are background-corrected, and standard deviations (k = 1) are listed for each scan.

| Activity (Bq) | iQID |  |
| --- | --- | --- |
|  | Count rate (s<sup>-1</sup>) | Standard deviation |
| 291.81 | 186.80 | 0.54 |
| 138.89 | 97.09 | 0.96 |
| 67.27 | 53.70 | 0.32 |
| 28.90 | 21.91 | 0.23 |
| 27.89 | 21.31 | 0.18 |
| 14.54 | 10.86 | 0.23 |
| 2.705 | 2.134 | 0.053 |
| 0.000 | 0.000 | - |

#### Cross-modality multi-scale phantom benchmarking

The initial activity of a ^177^Lu solution in an Eppendorf tube was (5.90 ± 0.26) MBq, measured with an HPGe detector. The combined standard uncertainty (4.4%) accounts for the detector calibration accuracy (3.0%) (33) and the spectral measurement consistency (3.2%), calculated as the standard deviation between activity estimates from the 113 keV and 208 keV photopeaks of (5.90 ± 0.19) MBq. Following dilution and decay, the final reference activities at each imaging time point were (0.200 ± 0.010) MBq in 0.850 mL injected into the Derenzo phantom and (0.197 ± 0.015) MBq in 0.213 mL dispensed onto the SML phantom. These estimates incorporate uncertainties from the HPGe characterization, decay correction, and the cumulative systematic error of the volumetric procedures. From μSPECT/CT, the total cumulative activity within each whole phantom volume was (0.174 ± 0.009) MBq (Derenzo) and (0.140 ± 0.007) MBq (SML), where the uncertainty (∼5%) reflects reported system cross-calibration accuracy (34,35). For iQID, the summed activity across all layers of SML phantom (whole-area per layer) was (0.184 ± 0.021) MBq, and the layer-wise activities were (19.5 ± 2.2) kBq, (19.7 ± 2.3) kBq, (19.4 ± 2.2) kBq, (19.8 ± 2.3) kBq, (16.8 ± 1.9) kBq, (18.1 ± 2.1) kBq, (16.7 ± 1.9) kBq, (18.1 ± 2.1) kBq, (18.9 ± 2.2) kBq, and (17.2 ± 2.0) kBq. The iQID activity uncertainty (11.4%) was dominated by the mean absolute relative residual of the calibration curve; by comparison, the relative count rate uncertainty from counting statistics and repeatability was < 0.5% per layer (∼1.1 million counts) and was therefore negligible in the total uncertainty. All values were decay-corrected to their respective μSPECT/CT scan times. To account for the geometric efficiency difference between the 0.1 mm well depth and the flat calibration source, a geometric correction factor (GF) of 0.95 derived from Geant4 Monte Carlo simulations was applied, with negligible associated simulation uncertainty (< 0.03%). After correction, the final iQID-derived total activity was (0.194 ± 0.022) MBq. Representative images are shown in Figure 6, and quantitative comparison results are summarized in **Table 6**.

**Figure 6.**
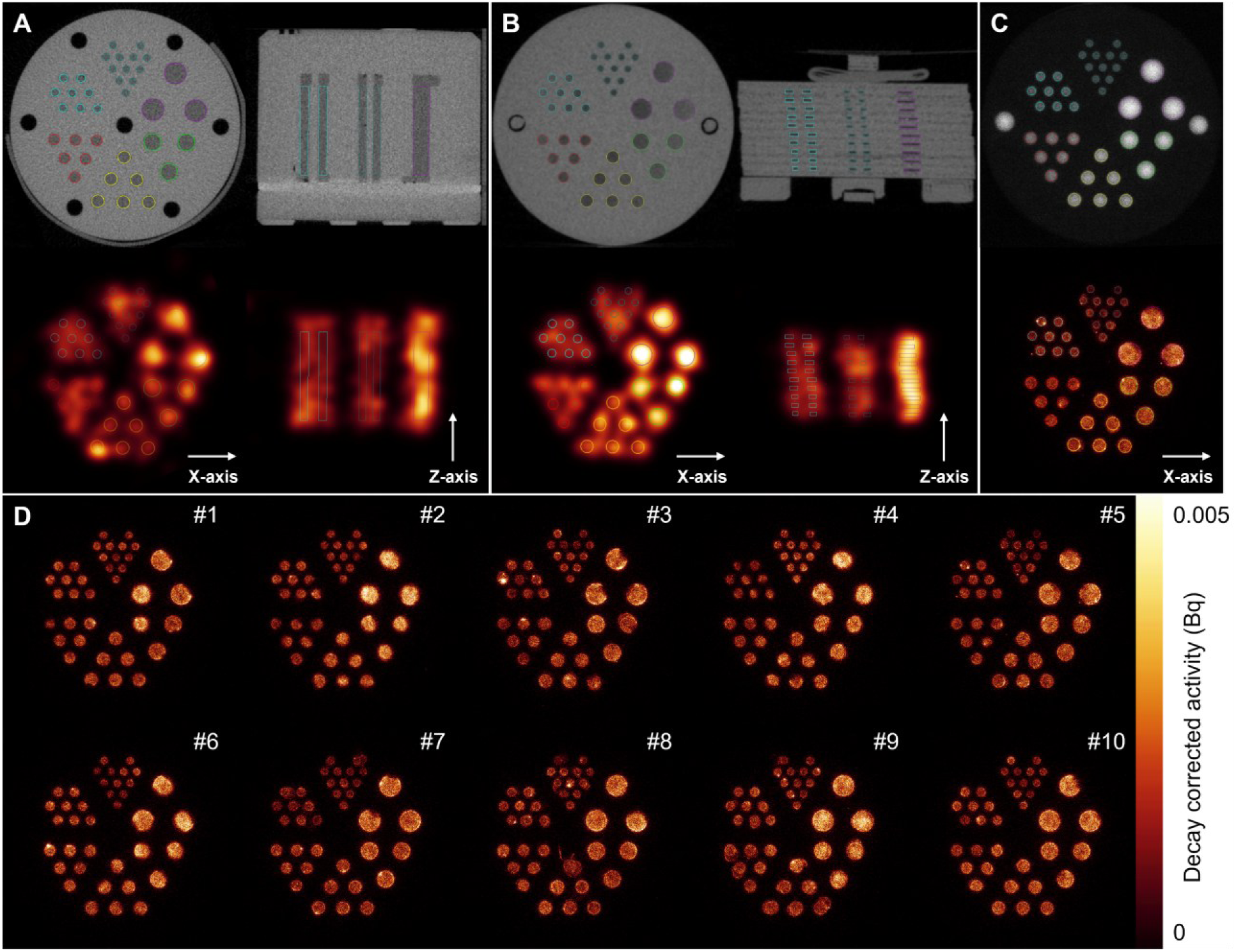
Quantitative ^177^Lu activity distribution with ROI delineations from 0.8 mm to 2.5 mm diameter wells. (A) Derenzo and (B) SML phantoms imaged with μSPECT/CT, showing ROI placement over CT images (top row; axial and coronal views) and co-registered activity distributions (bottom row). (C) iQID shadowgraph used for ROI delineation (top) and a representative planar autoradiography image of a single-layer SML phantom (bottom). (D) Planar iQID images of individual layers. The color scale applies to all activity images.

**Table 6.** Cumulative ^177^Lu activity in Derenzo and SML phantoms measured by μSPECT/CT and iQID. Percent differences are relative to injected activities. iQID_GF_ denotes results with geometric correction applied.

| Phantom |  | Total Activity (MBq) | %Difference |
| --- | --- | --- | --- |
| Derenzo | Injection | $0.200 \pm 0.010$ | - |
| | $\mu\text{SPECT}$ | $0.174 \pm 0.009$ | -13.1 |
| SML | Injection | $0.197 \pm 0.015$ | - |
| | $\mu\text{SPECT}$ | $0.140 \pm 0.007$ | -28.7 |
| | iQID | $0.184 \pm 0.021$ | -6.3 |
| | iQID<sub>GF</sub> | $0.194 \pm 0.022$ | -1.4 |

#### Recovery coefficient analysis

To evaluate partial volume effects in small-scale imaging, ROIs were delineated individually for each 10 mm length rod in the Derenzo phantom and each 0.1 mm depth well in the SML phantom (Figure 6A**–C**). The reference activity concentrations, derived from HPGe quantification and micropipette dispensed volumes, were (0.236 ± 0.012) MBq/mL for the Derenzo and (0.924 ± 0.068) MBq/mL for the SML phantom. For μSPECT imaging of the Derenzo phantom, activity concentrations ranged from (0.109 ± 0.005) MBq/mL (2.5 mm) to (0.047 ± 0.002) MBq/mL (0.8 mm), corresponding to relative differences of –53.8% to –80.1%. Similarly, μSPECT measurements from the SML phantom ranged from (0.138 ± 0.007) to (0.048 ± 0.002) MBq/mL, reflecting –85.0% to –94.8% deviation from the reference. In contrast, iQID imaging of the SML phantom yielded activity concentrations from (0.660 ± 0.075) to (0.351 ± 0.040) MBq/mL (–28.5% to –62.0%). After applying the geometric correction factor (GF = 0.95), the adjusted iQID values ranged from (0.743 ± 0.085) to (0.466 ± 0.053) MBq/mL (–19.5% to –49.5%). RCs calculated from these measurements are summarized in **Table 7**. The corresponding individual rod or well profile graphs of activity concentration on the X- and Z-axes are provided in Figure 7D**–F**. The voxel dimensions were 0.02 mm for iQID and 0.08 mm for µSPECT/CT.

**Figure 7.**
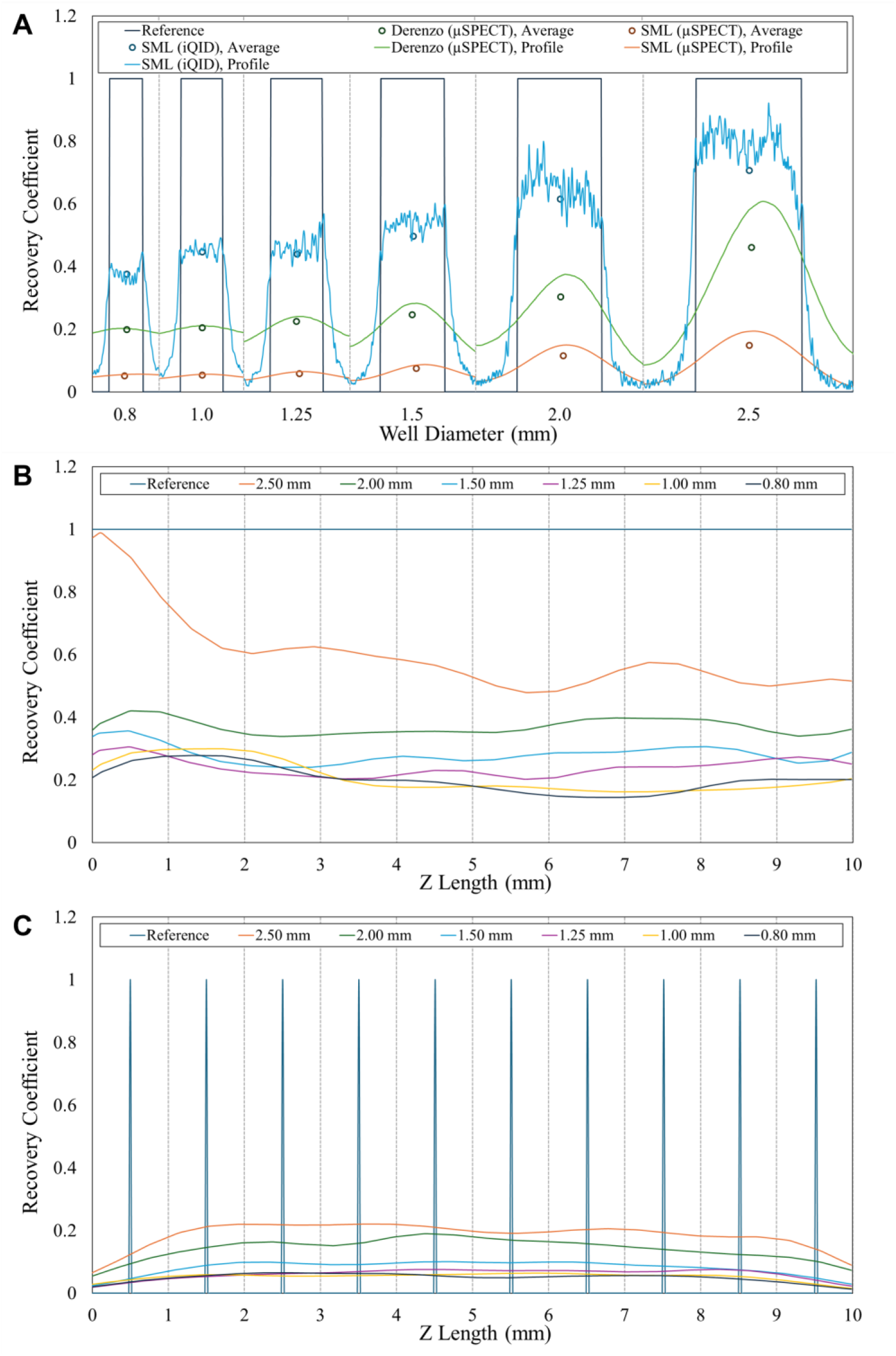
Recovery coefficient (RC) analysis. (A) X-axis RC profiles against each rod/well diameter for both phantoms and modalities, compared to the ideal reference profile. (B) Derenzo and (C) SML Z-axis RC profiles measured by μSPECT across different rod/well diameters.

**Table 7.**
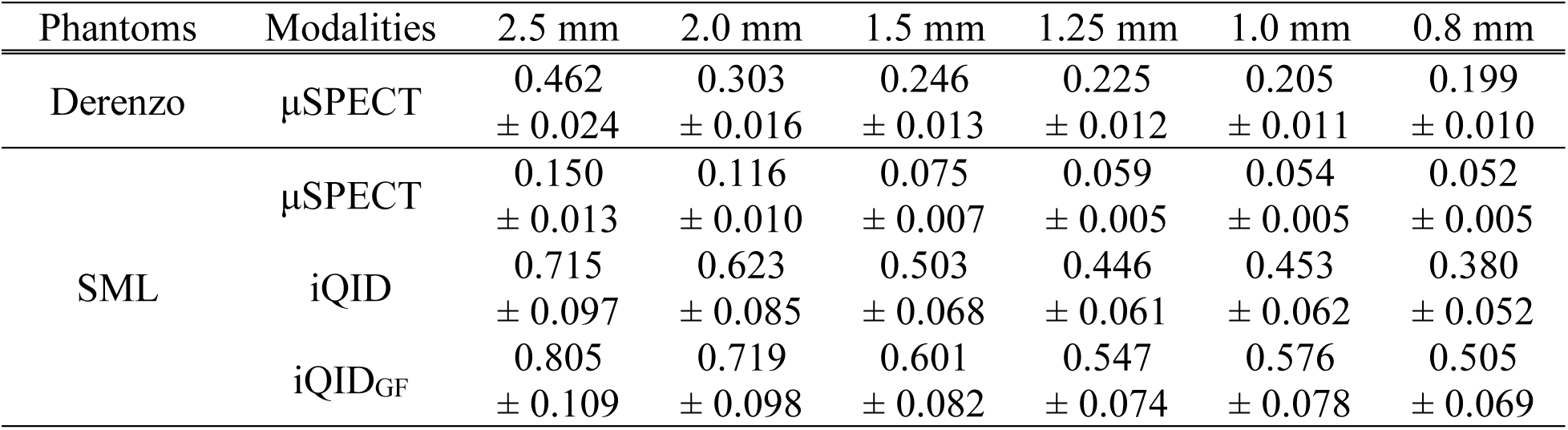
Recovery coefficients (RCs) for rod/well diameters derived from μSPECT/CT (Derenzo, SML) and iQID (SML) images. iQID results are shown with and without geometry correction.

## DISCUSSION

This study demonstrates a quantitative approach for mapping activity distributions of the β-emitter ^177^Lu for applications in small-scale preclinical RPT research using the iQID digital autoradiography system. Characterization confirmed the system’s high-resolution performance, with an intrinsic resolution of ∼20 μm and an effective resolution of ∼43 μm for ^177^Lu. To reflect preclinical *ex vivo* measurements of thin tissue sections, the characterization framework accounted for geometric and acquisition conditions relevant to such specimens by measuring and subtracting background, determining the MDA for ^177^Lu, and evaluating signal dependence on sample depth. Guided by these results, calibration measurements were then performed using acquisition settings optimized for preclinical tissue imaging, yielding accurate activity quantification. While cross-modality multi-scale phantom benchmarking showed expected quantitative accuracy for μSPECT imaging (−13.1% for Derenzo and −28.7% for SML), the iQID system demonstrated superior quantitative accuracy for microscale activity (−1.4%) and activity concentrations with high RCs (0.5−0.8), effectively resolving small structures where μSPECT performance was limited by partial volume effects.

Our work extends the scope of previous digital autoradiography studies (11,13,21), predominantly focused on short-range α-emitters, to the β-emitting theranostic radionuclide ^177^Lu. We established a rigorous characterization framework that integrates Geant4 Monte Carlo simulation to physically interpret the more complex detector response associated with continuous energy β spectra. Beyond standard source calibration, we addressed the complexities of realistic tissue measurement scenarios by quantifying detection efficiency and depth-dependent signal modulation. Furthermore, the cross-modality validation against µSPECT/CT provides a critical bridge between macroscopic and microscopic imaging modalities. By analyzing RCs, we confirmed the capacity of iQID to resolve microscale activity distributions and concentrations that are obscured by partial volume effects in µSPECT, thereby laying the groundwork for high-fidelity multi-scale dosimetry in future *in vivo* and *ex vivo* applications.

This study has limitations related to the physical properties of β-emitters and the optical characteristics of the detector. First, localizing β-emitters is inherently more complex than ⍺-emitters due to the longer particle range and increased scattering, which generates light in the phosphor screen further from the source origin. While capillary filters can mitigate this lateral spreading (36), they reduce total light transmission, necessitating longer scan times and potentially introducing geometric uncertainties. Furthermore, pipette-based dispensing is susceptible to stochastic drying dynamics, notably the coffee-ring effect, which can concentrate activity at droplet boundaries. This microscopic heterogeneity can create localized regions of high flux density, which can increase susceptibility to pile-up in iQID and exacerbate partial volume and spill-out effects in µSPECT/CT. Second, the iQID optical chain is sensitive to ambient light. In our setup, exposure of the Gadox screen to room light increased the measured background rate by approximately threefold. To standardize measurements, light exposure was minimized and acquisitions were initiated only after the background returned to baseline, which typically required ∼30 minutes after any substantial room light exposure. Finally, high activity samples introduced pile-up effects, in which temporally coincident scintillation events merge within a single frame in confined source regions, leading to count loss and reduced apparent count rates at high local flux density. Under the calibration geometry used in this study (three 10 µL drops), this behavior became observable at activities above approximately 70 Bq. Consequently, detector settings such as input voltage should be optimized to balance background noise against signal detection, and calibration curves should be specific to the activity range being measured.

The cross-modality benchmarking study also presented experimental limitations, particularly regarding the SML phantom preparation. The ^177^Lu solution was manually dispensed into the wells, which resulted in non-uniform drying patterns and minor activity deposition outside the wells. These inconsistencies likely biased the geometric correction factors (which assumed uniform, perfect discs) and explain anomalies such as the higher RC observed for the 1.0 mm well compared to the 1.25 mm well using iQID. To mitigate these challenges, future ^177^Lu phantom preparations could leverage metrology-grade inkjet deposition systems to generate highly uniform, well-characterized activity distributions within each well or on the surface (37,38). Moreover, the comparison highlighted the intrinsic spatial resolution limits of µSPECT/CT (the lowest of 0.4 mm resolution) for this application. While µSPECT provided approximate estimates of total activity (–28.7% relative to the reference), it substantially underestimated activity concentrations within the millimeter-scale wells (–91.6% on average).

This severe bias is consistent with partial volume effects inherent to 2D surface source geometries, where the source layer thickness is significantly smaller than the system’s axial resolution, leading to substantial spill-out effects in the reconstructed volume (39). Notably, the SML phantom design intentionally creates a quasi-continuous 10 mm length rod appearance to μSPECT/CT while enabling post-imaging disassembly for layer-resolved autoradiography, providing the cross-modality test object capable of validating layer-specific activity distribution (Figure 7).

Overall, this work establishes a comprehensive framework for characterizing and calibrating the iQID system for ^177^Lu, addressing a critical need in small-scale RPT dosimetry. Our thin-layer phantom results demonstrate that this platform delivers quantitative accuracy and spatial resolution significantly superior to preclinical µSPECT/CT, supporting its potential for precise activity mapping in tissue sections. In a preclinical setting, this enables a multi-scale activity mapping for theranostic radionuclides. By registering macroscopic μSPECT/CT to track ROI retention with microscopic iQID autoradiography to capture intra-ROI heterogeneity, providing the direct activity inputs required for Monte Carlo simulation–based internal dosimetry platforms (40) to estimate microscale absorbed dose distributions. These microscale dosimetric data can subsequently be co-registered with biological readouts, such as hematoxylin and eosin (H&E) or phosphorylated histone H2AX (γH2AX) staining, to correlate dose with biological effect at the cellular level (17,18). Furthermore, the detector’s particle- and energy-dependent response offers the potential to discriminate spatial distributions in multi-radionuclide cocktail studies or to assess progeny redistribution in α-particle therapies (e.g., ^225^Ac and ^213^Bi accumulations in kidneys) (20,41). Realizing the full clinical potential of this multi-scale approach will require future integration of image registration techniques, derivation of subject-specific pharmacokinetic parameters from in vivo imaging, and the optimization of clinical biopsy protocols.

## CONCLUSION

This study presents the characterization and calibration of the iQID digital autoradiography system for direct quantitative imaging of the β-emitter ^177^Lu. Using surface-dispensed sources, we established a rigorous protocol for converting count rates to activity and evaluated the detector’s physical response. The cross-modality benchmarking comparison with μSPECT/CT validated this quantitative framework using the layered phantoms, confirming the iQID’s superior capability to resolve microscale activity concentrations that are otherwise obscured by partial volume effects in standard nuclear medicine imaging. These results provide a practical basis for applying iQID to activity estimation in thin tissue sections and registration with μSPECT, enabling high-resolution inputs for multi-scale RPT dosimetry research.

## DECLARATIONS

**Ethics approval and consent to participate:** Not applicable.

**Patient consent for publication:** Not applicable.

**Availability of data and material:** All data relevant to the study are included in the article.

**Competing interests:** BPB is the Co-Founder and Scientific Advisor of Voximetry, Inc.

## Acknowledgements

The authors’ work is supported in part by grants from NIH NCI P01CA250972. The authors would like to thank the University of Wisconsin Carbone Cancer Center (UWCCC) and Small Animal Imaging & Radiotherapy Facility for supporting this project.

## Authors’ contributions

OK performed the primary experimental and simulation work, data curation, and analysis; OK and SPJ conducted secondary experiments for characterization. AOA managed iQID and JJJ managed μSPECT/CT imaging modalities. JBS designed and fabricated 3D-printed components. LEW and MB prepared and processed radionuclide solutions. EA managed HPGe detector. BWM and DEB provided technical guidance on digital autoradiography and study design. RH, LAD, and BPB provided laboratory infrastructure and experimental resources. BPB led study conception, function acquisition, project administration, supervision, and data interpretation. OK and BPB drafted the manuscript; all authors reviewed and approved the final version.

## Competing interests

JJJ is cofounder and Chief Executive Officer of RPT Labworks and cofounder and Chief Science Officer of Phantech. BWM is founder and president of Qsint. RTH is cofounder and Chief Science Officer of RPT Labworks and a member of scientific advisory board for Archeus Technologies. LAD is cofounder and Chief Science Officer of Standard Imaging, Inc. BPB is cofounder and scientific advisory board for Voximetry, Inc. All other authors declare they have no competing interests.

**Correspondence** and requests for materials should be addressed to Bryan P. Bednarz.

## Footnotes

1 Certain commercial equipment, instruments, or materials are identified in this paper to foster understanding. Such identification does not imply recommendation by the National Institute of Standards and Technology, nor does it imply that the materials or equipment identified are necessarily the best available for the purpose.

